# *C. difficile*-associated antibiotics prime the host for infection by a microbiome-independent mechanism

**DOI:** 10.1101/728170

**Authors:** Jemila C. Kester, Douglas K. Brubaker, Jason Velazquez, Charles Wright, Douglas A. Lauffenburger, Linda G. Griffith

## Abstract

The most clinically relevant risk factor for *Clostridioides difficile*-associated disease (CDAD) is recent antibiotic treatment. Though most broad-spectrum antibiotics significantly disrupt the structure of the gut microbiota, only particular ones increase CDAD risk, suggesting additional factors might increase the risk from certain antibiotics. Here we show that commensal-independent effects of antibiotics collectively prime an *in vitro* germ-free human gut for CDAD. We found a marked loss of mucosal barrier and immune function with CDAD-associated antibiotic pretreatment distinct from pretreatment with an antibiotic unassociated with CDAD, which did not reduce innate immune or mucosal barrier functions. Importantly, pretreatment with CDAD-associated antibiotics sensitized mucosal barriers to *C. difficile* toxin activity in primary cell-derived enteroid monolayers. These data implicate commensal-independent host changes in the increased risk of CDAD with specific antibiotics. Our findings are contrary to the previously held belief that antibiotics allow for CDAD solely through disruption of the microbiome. We anticipate this work to suggest potential avenues of research for host-directed treatment and preventive therapies for CDAD, and to impact human tissue culturing protocols.

## Introduction

*Clostridioides difficile*-associated disease (CDAD) is a CDC urgent public health threat^1^, with 453,000 incident cases in the U.S. in 2011^2^. CDAD is estimated to account for more than 44,500 deaths and over $5 billion in related healthcare costs in the United States each year^3^. CDAD treatment failure is increasing due to rising levels of antibiotic resistant and hypervirulent strains of *C. difficile* (reviewed in^4^) and high rates of persistent and recurrent infections (reviewed in^5^). New treatment and prevention strategies are needed. A promising strategy is host directed therapy^6^; yet, this requires a better understanding of how a person becomes susceptible to CDAD.

CDAD pathogenesis requires the outgrowth of the etiologic agent, *Clostridioides difficile* (*C. difficile*), in the gastrointestinal tract. While a functional gut microbiome is able to prevent the outgrowth of *C. difficile*^7^, in large part due to bacterial-dependent production of secondary bile acids^8,9^, loss of a functional gut microbiome allows for outgrowth of the colony. Once quorum is reached, the bacteria begin secreting toxins^8^, specifically TcdB, which is primarily responsible for the disease’s symptoms and pathogenesis^9^.

The most clinically relevant risk factor for CDAD is recent antibiotic treatment^10^. While there is substantial evidence supporting a causal link between microbiome disruption by antibiotics and CDAD (reviewed in^11^), the hypothesis that recent antibiotic treatment is the sole causal mechanism for CDAD explains neither the rising rates of CDAD—independent of recent antibiotic treatment^12^—nor the observation that nearly half of all community acquired cases present without prior antibiotic exposure^13^. Furthermore, antibiotic treatment alone produces fewer and less consistent differences in microbial community structure than are observed clinically between patients with and without CDAD^14^. Notwithstanding the frequency of proteobacteria blooms following antibiotic exposure^15^, no shared taxonomic change has been identified in successful fecal microbial transplant donors^16^ or recipients^17^, and alterations to bacterial load do not correlate with risk of CDAD^18^. Taken together, these data suggest a previously overlooked commensal-independent host contribution to antibiotic-associated risk of CDAD.

Recent work has shown that host-acting drugs have a significant effect on bacteria^19^. The inverse has also been shown: a commonly prescribed antibiotic cocktail alters the mitochondrial function of enterocytes in germ-free mice^20^, demonstrating the commensal-independent effect of anti-bacterials on the host in this rodent model. Yet, the effects of CDAD-associated antibiotics on the host—especially the *human* host—and how these effects might contribute to CDAD is not known. Though previous studies have explored the effects of antibiotics in general on the host in a variety of animal models, isolating the effects of particular antibiotics on host-dependent mechanisms of antibiotic-associated CDAD requires a controlled study assessing these mechanisms for multiple antibiotics with varying degrees of CDAD-associated risk in the same experimental context, preferably with models that include elements of the human mucosal barrier response. It is only in this context of a multi-factorial study design that the most translationally relevant biological findings can be identified and validated in primary human donor tissues.

## Results

### CDAD-associated antibiotics induce distinct changes to host gene expression

To test the commensal-independent effects of antibiotics on the host, we used a transwell-based *in vitro* epithelial barrier without bacteria to model a germ-free human gut^21,22^. We treated mature mucosal barriers with antibiotics with CDAD odds ratios, from one (no risk) to 17 (highest risk; Supplemental Table 1) in order to achieve complete coverage of the CDAD risk landscape. We dosed from the basal side to mimic intravenous administration due to its increased risk of CDAD^23^, using clinically-relevant dose ranges. Tigecycline is an intravenous tetracycline derivative that does not increase the risk of CDAD^24^. We used clindamycin and ciprofloxacin for CDAD-associated antibiotics as they have the highest risk of CDAD^24,25^. Tigecycline and clindamycin share a similar mechanism of action, both targeting bacterial translation machinery. Conversely, ciprofloxacin inhibits bacterial DNA replication. All three are considered broad-spectrum antibiotics.

After 24 hours of exposure, RNA-seq identified gene expression changes under high and low treatment conditions, with the largest number of transcriptional changes being driven by ciprofloxacin treatment (Fig 1B-E). We found DMSO treatment at high concentration had significant effect on gene expression, and therefore performed subsequent analysis using the low concentrations of both DMSO and tigecycline (Supplemental Fig S1).

**Fig 1:**
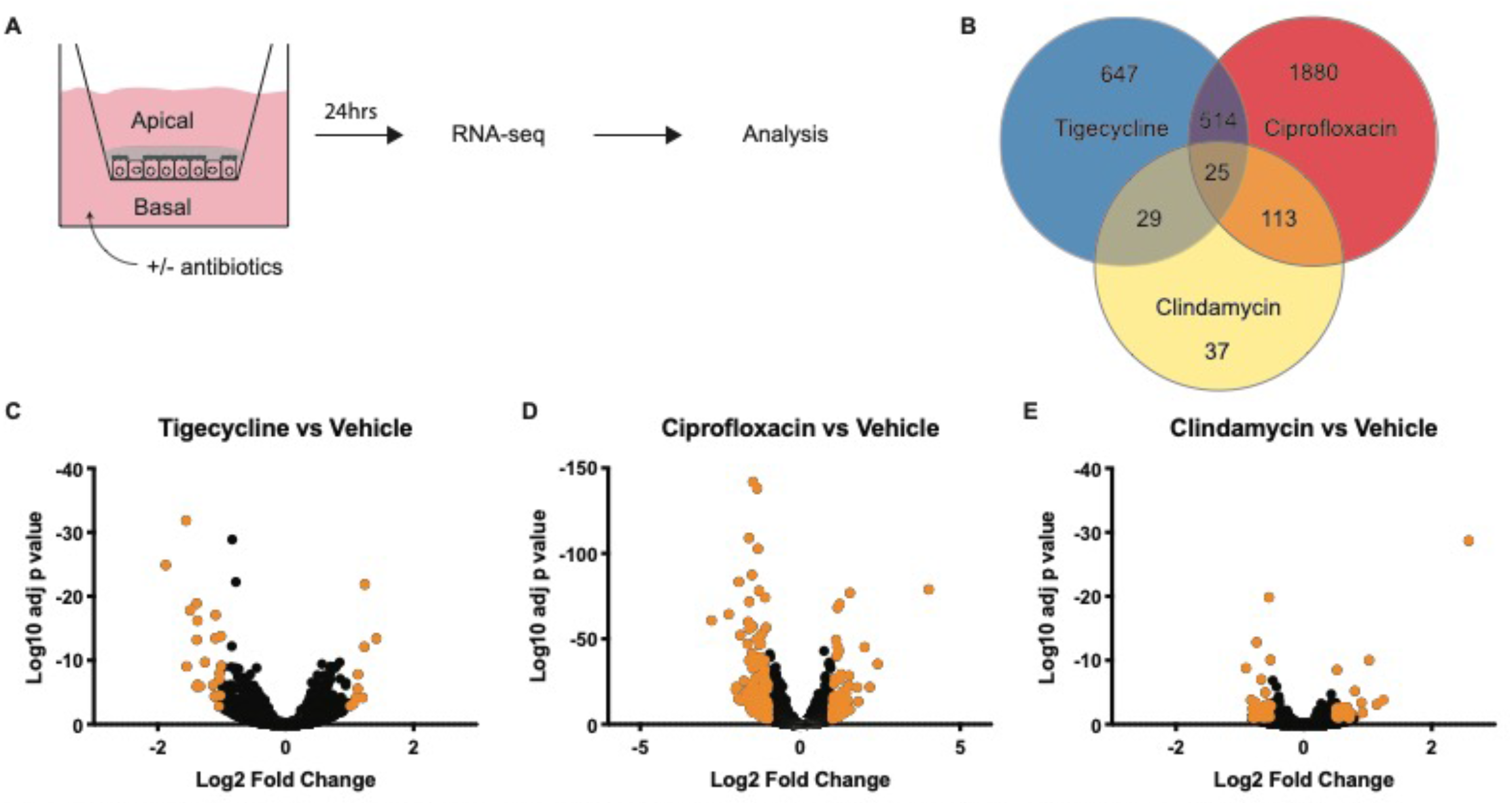
RNA-seq identifies differential gene expression changes following antibiotic treatment. A) Schematic of RNA-seq procedure. 6 transwells per condition. B) Venn diagram and color key of gene expression patterns. C-E) Volcano plots of gene expression changes for indicated antibiotic over vehicle. Highlighted points have at least a 2-fold change with an adjusted p value of <0.01.

Unsupervised hierarchical clustering of the 606 genes with statistically significant gene expression changes in at least two treatment groups revealed antibiotic-specific alterations of the gut transcriptome (Supplemental Fig S2). This clustering showed varying patterns of transcriptional response to treatment: several transcripts shared similar expression configurations across experimental conditions, some had dose-dependent effects correlating to increasing or decreasing CDAD risk, still others exhibited more complex behaviors not apparent from the initial clustering. However, the ciprofloxacin-driven expression changes dominated the clustering, highlighting the need for more nuanced computational analysis.

In order to identify genes with transcriptional changes shared between both CDAD-associated antibiotics, we used a self-organizing map (SOM). A SOM is a neural network-based unsupervised clustering technique that groups similar observations together on the SOM neurons. Here, we used the SOM to cluster gene transcript fold changes across antibiotic treatments to identify genes with expression changes associated with CDAD risk. Similar to other dimensionality reduction techniques, such as principal components analysis (PCA), SOMs produce a low-dimensional projection of high-dimensional data that facilitates visualization of patterns. However, unlike PCA or our previous hierarchical clustering, the SOM analysis merges two important features simultaneously: (i) it incorporates information about the expected number of clusters in the data by defining the number of SOM neurons based on experimental design (number of conditions); and (ii) it allows the data to drive identification of the most informative groups among those clusters (i.e., SOM neurons).

The architecture of the SOM employed here to map the 606 significant genes is based on increased or decreased gene expression (2 directions) in each of three (3) experimental conditions, with an extra neuron for noisy profiles (2^3^+1=9 neurons). Genes with similar expression patterns cluster in a node, with the number of genes per node indicated (Fig. 2A). Plotting neighbor weight distances allows for the visualization of similarities between nodes (Fig. 2B).

**Fig 2:**
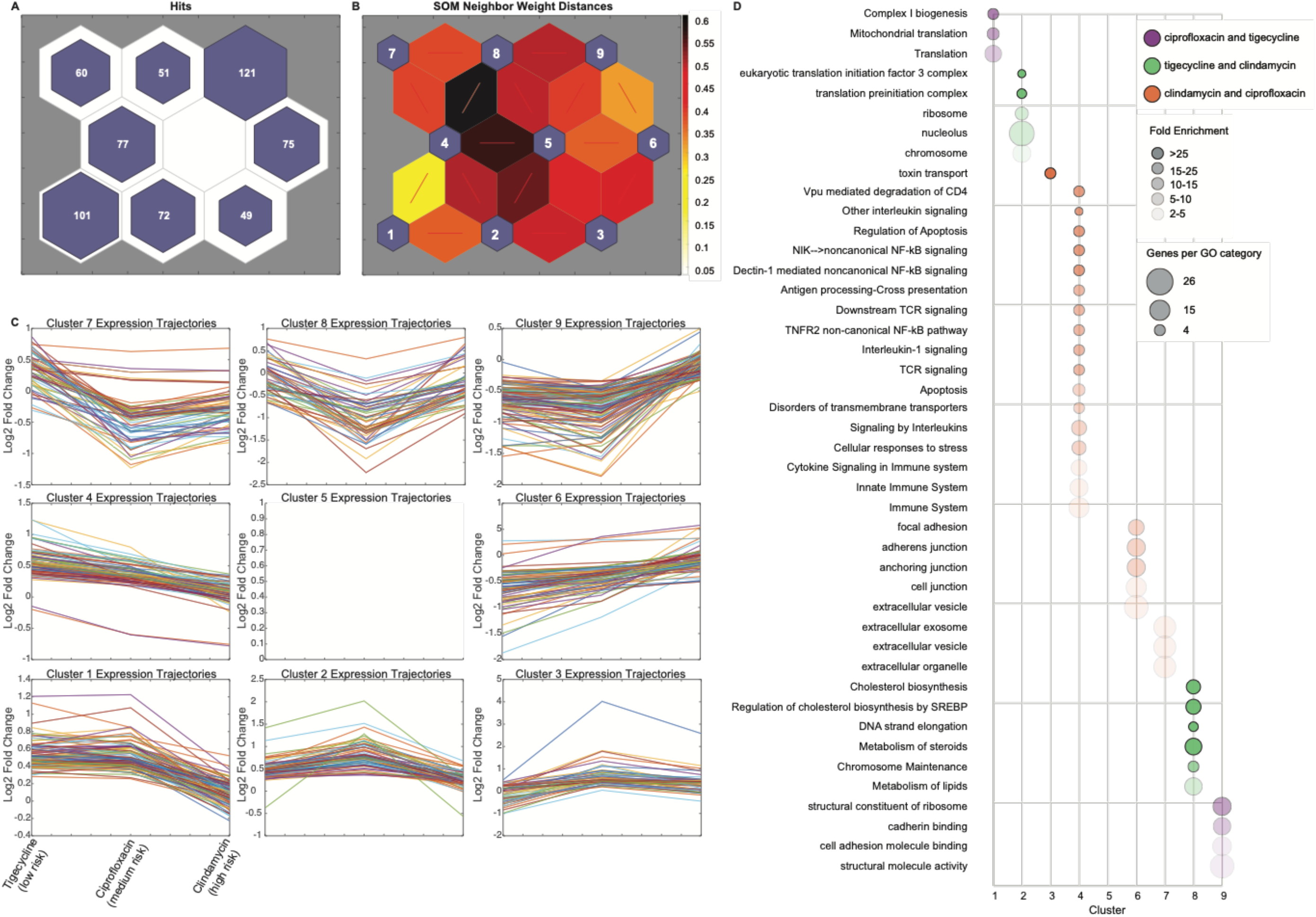
Self-organizing map (SOM) predicts CDAD-associated antibiotics may alter barrier and immune functions. A) SOM of 606 genes with statistically significant expression changes by RNAseq. B) SOM neighbor weight distances indicate level of similarity between each node pair. C) Line graphs of all genes in each node, plotted as increasing risk of CDAD on the x-axis by standardized fold change on the y-axis. D) Bubble chart showing overrepresented GO terms for clusters as indicated using PANTHER overrepresentalion test. FDR <0.05.

Each neuron of the SOM captured gene expression responses to antibiotic treatment that grouped according to changing CDAD risk ratios. These patterns could then be investigated by plotting line graphs of the gene fold changes across increasing CDAD risk for each node (Fig. 2C). Two nodes identified gene expression responses that were specifically elevated (Node 3) or repressed (Node 7) in response to CDAD-associated antibiotic treatment. Another two nodes (4 and 6) captured risk ratio-dependent changes in gene expression responses to CDAD-associated antibiotic treatment, with genes on Node 4 being more downregulated and genes on Node 6 being more upregulated in antibiotics with higher CDAD risk ratios. Altogether, nodes 3, 4, 6, and 7 capture a set of 261 genes with expression patterns common among ciprofloxacin and clindamycin that indicated a shared pattern of expression unique to the CDAD-associated antibiotics (Fig. 2C), despite different mechanisms of action between ciprofloxacin and clindamycin, and tigecycline and clindamycin being similar.

In order to identify the biological functions associated with CDAD-associated antibiotic treatment, we performed Gene Ontology Enrichment Analysis (GOEA) of each node (Fig. 2D, Supplemental Table 2). We would expect nodes that cluster by mechanism of action to be enriched in related GO terms. For instance, the gene expression responses common to tigecycline and clindamycin (Nodes 2 and 8) were enriched for the cellular targets of those drugs, translation machinery and chromosome maintenance (Fig. 2D). It is important to note that these targets are considered bacterial cellular components, yet we found they impacted mammalian cells. This finding from the SOM clustering that grouped known target-associated gene expression responses to tigecycline and clindamycin provided an important positive control for interpreting the biological functions associated with the other SOM neurons.

We then analyzed the SOM clusters that captured genes with shared expression response patterns to CDAD-associated antibiotics (Nodes 3, 4, 6, and 7), and distinct from low risk, to generate mechanistic hypotheses of host-dependent mechanisms of CDAD. The GOEA functional annotations of CDAD-associated antibiotic treatment showed an accumulation of cellular toxins in the cell via retrograde secretion (Node 3: toxin transport) coupled with a decrease in secretion out of the cell (Node 7). We found that as antibiotic CDAD risk ratios increased, genes associated with immune signaling GO terms were suppressed (Node 4) and genes associated with cell-cell and cell-ECM connections were increased (Nodes 6 and 7) in a dose-dependent manner. Overall, GOEA of these SOM suggested that treatment with CDAD-associated antibiotics resulted in alterations to transport of extracellular components out of the cell, toxins into the cell, and a reduced immune capacity after only 24hrs of treatment.

### CDAD-associated antibiotics reduce mucosal barrier and immune functions

Based on the results of the SOM analysis, we hypothesized that CDAD-associated antibiotic treatment would result in acute effects of impaired epithelial barrier and innate immune cell function and that these would be present after chronic exposure as with *in vivo* antibiotic treatment patterns. We tested these SOM predictions experimentally using three complementary levels of *in vitro* models: (i) acute effects (24 hr) on epithelial barrier function; (ii) chronic (3 days) effects on epithelial exposure to toxin; and (iii) acute effects on innate immune cell function. We assessed the barrier function of mucosal barriers following antibiotic treatment. We find increased death of cells in the monolayers with ciprofloxacin treatment, which agrees with previous work using significantly higher concentrations^26^, and clindamycin, which has not previously been demonstrated (Supplemental Fig S3). Despite this, none of the antibiotics used affected the physical integrity of the barrier as determined by transepithelial electrical resistance 24 hours post treatment (Supplemental Fig S4).

To better mimic the 3-7 day course of antibiotics in routine *in vivo* human treatment patterns, we extended the treatment period to a 3 day basal dose. We quantified both mucin gene expression and secretion as they strongly influence microbial interactions with the mucosal barrier. Total cell-bound (Fig. 3A) mucin was reduced with CDAD-associated antibiotics, while mucins in low risk CDAD treatment groups remain unchanged (Fig. 3A, B). Secreted mucins were reduced with both CDAD-associate antibiotics, though the clindamycin treatment group does not reach statistical significance (Fig 3B). This is recapitulated in primary cell-derived 2D enteroids: one of the main membrane-bound mucins in the colon^27^, *muc17*, is reduced with ciprofloxacin but not clindamycin or tigecycline treatment (Fig. 3C), suggesting another mucin is altered with clindamycin treatment to account for the loss of total cell-bound mucins.

**Fig 3:**
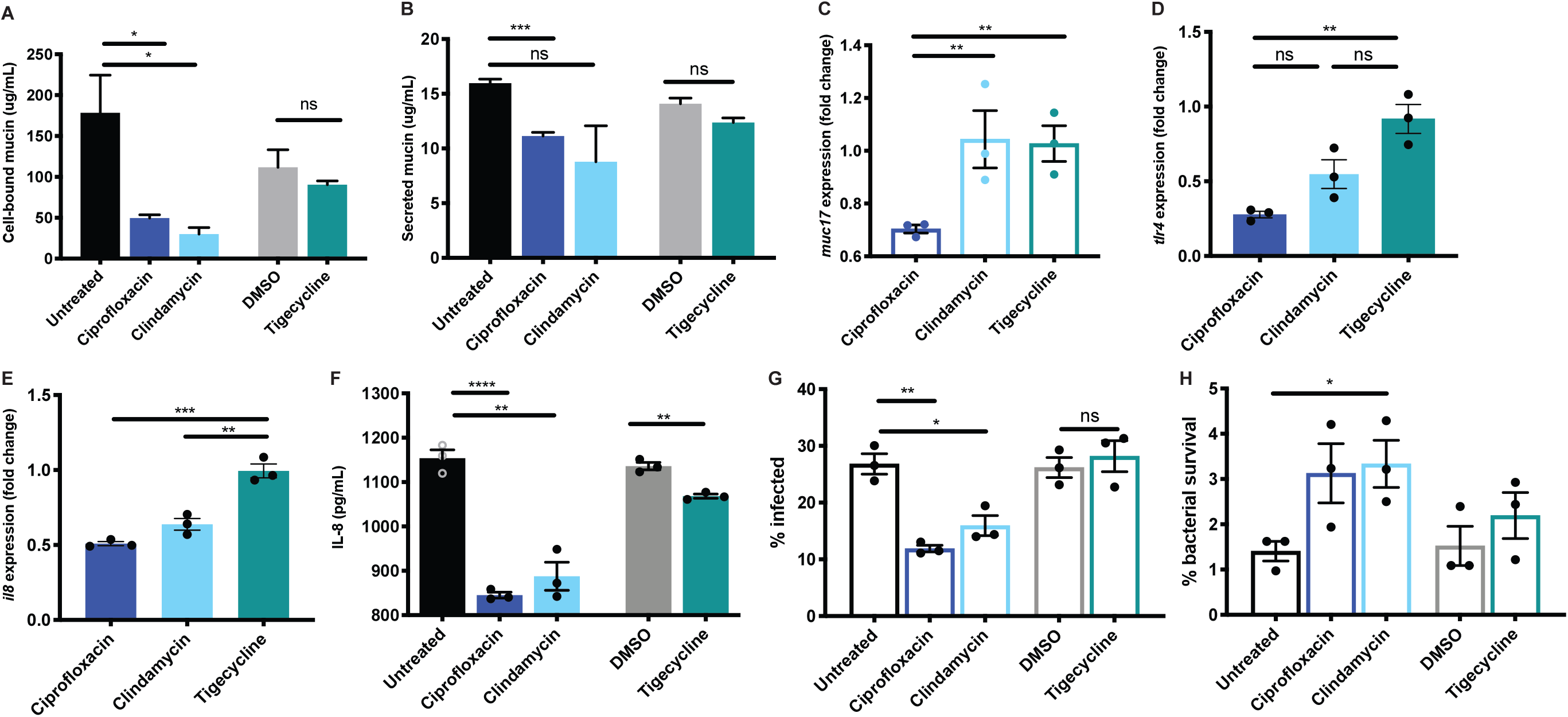
CDAD-associated antibiotics reduce mucosal barrier and immune functions. A) Cell-bound and B) secreted mucin quantification. C) Expression fold changes of *muc17* in primary cell-derived 2D enteroids relative to vehicle by qPCR. D-F) immune-competent monolayers: D) *tlr4* and E) *il8* gene expression fold change with LPS stimulation relative to vehicle by qPCR. F) Total IL-8 basal secretion with LPS stimulation by ELISA. Representative data, experiments repeated 2-3 times with similar results, n=3 per experiment. G) Quantification of phagocytosis and H) intracellular killing of *E. coli* by primary macrophages pre-treated for 3 days with indicated antibiotic. Primary cell data in open bars, cell line data in filled bars. Representative data, experiments repeated 2-3 times with similar results, n=3 per experiment. Statistical significance determined by student’s t test; *<0.05, **<0.01, ***<0.001, ****<0.0001.

To assess the effect of extended, low-dose antibiotic treatment on immune function, we treated an immune-competent mucosal barrier for 3 days with each antibiotic, again dosing from the basal side. IL-8 secretion is the primary chemokine implicated in CDAD^28^. IL-8 is required for neutrophil recruitment to contain the infection, yet neutrophils are also implicated in progression of disease^29^. Thus, a delicate control over dissemination and clearance of neutrophils is likely required for resolution of infection.

We therefore assessed the effect of antibiotics on the ability of immune-competent mucosal barriers to induce *il8* expression and IL-8 secretion following LPS stimulation. LPS signals through TLR4 and *tlr4* gene expression should increase following its activation, yet *tlr4* expression did not increase with LPS stimulation following ciprofloxacin treatment (Fig 3D). Clindamycin treated barriers had lower levels of *tlr4* relative to vehicle, though this was not significantly lower than for tigecycline by student’s t test (Fig 3D).We found *il8* gene expression (Fig. 3E) is reduced following ciprofloxacin and clindamycin treatment but unchanged with tigecycline in LPS-treated barriers. IL-8 secretion (Fig. 3F) was reduced to a statistically significance extent in all treatment groups. It is likely the magnitude of change is important in the case of CDAD-associated antibiotics.

To test whether the immune cells are impaired in function, we performed phagocytosis and killing assays using GFP+ *E. coli*. We find that pre-treating macrophages with CDAD-associated antibiotics reduce both phagocytosis of *E. coli* (Fig. 4A) and subsequent killing of phagocytosed *E. coli* (Fig. 4B). Together, these data confirm loss of immune responsiveness with CDAD-associated antibiotics, which one can imagine might contribute to outgrowth of *C. difficile*.

**Fig 4:**
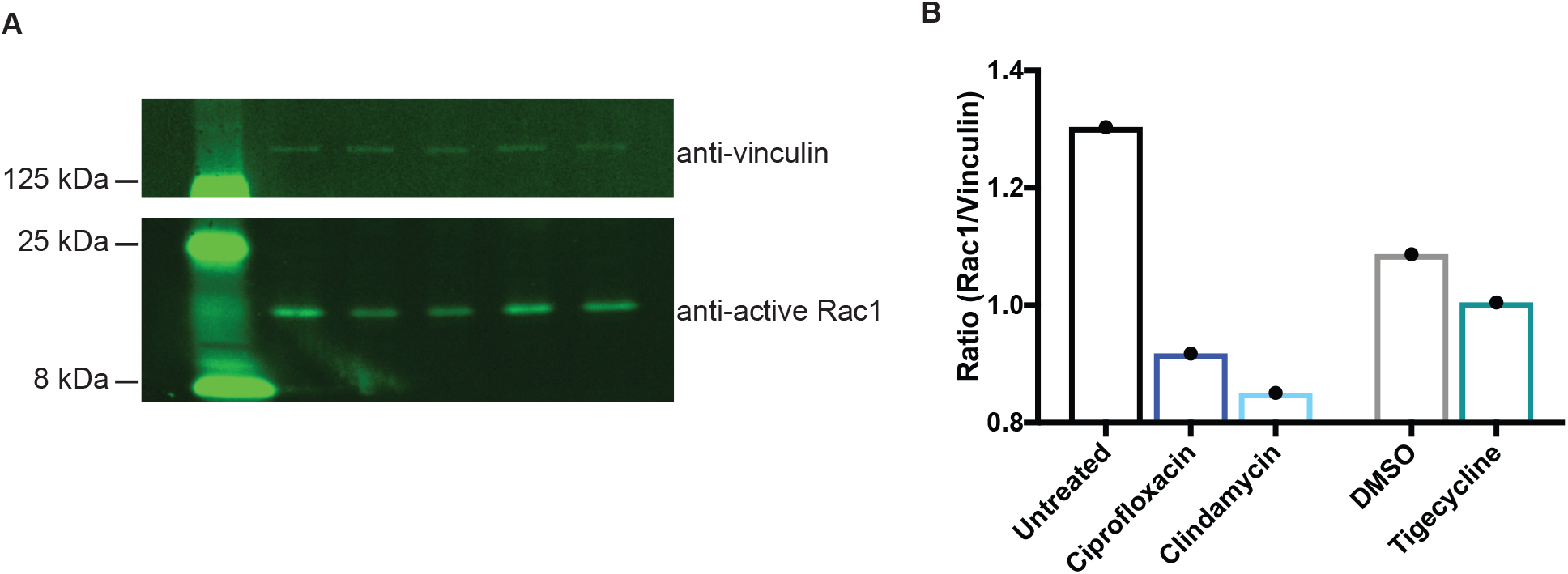
CDAD-associated antibiotic effect is recapitulated in primary tissue. A) Western blots show reduced active Rac1 in primary cell-derived monolayers. B) Quantification of western blot, Rac1:vinculin. Repeated twice with similar results, n=1 shown.

### Antibiotic effects are recapitulated in primary tissue

Previous work has shown the importance of mucus—primarily made up of mucins—on preventing *C. difficile* toxins from entering gut cell lines in culture^30^. CDAD’s pathology is driven by the cytotoxic effects of toxins, primarily TcdB^28^. TcdB enters epithelial cells through receptor-mediated endocytosis. Following acidification of the vacuole, the toxin enters the cytosol where it glucosylates its GTPase targets Rho, Rac, and Cdc42. This leads to actin depolymerization, characterized by visible rounding of the cell, and eventually to cell death. Our robust mucosal barrier is affected by TcdB as expected, by cell rounding and death as measured by holes in the monolayer after 48 hours of treatment with *C. difficile* filtered culture supernatant (Supplemental Fig S5).

To understand the translation of these altered barrier properties to potential impact in CDAD, we treated mucosal barriers with TcdB and measured its action by the loss of a cellular target, activated Rac-1, by western blot analysis. Both CDAD-associated antibiotics had deactivation of Rac1 at 24hrs while tigecycline and controls were still active by quantitative western blot (Fig. 4C, D), implicating a shared sensitization to *C. difficile* toxin from CDAD-associated antibiotics with a mechanism independent of commensals.

## Discussion

Here we demonstrate the convergent changes to the host from two separate CDAD-associated antibiotics in the absence of commensal bacteria. CDAD has been suggested to be a multi-phase system, where germination, outgrowth, and toxin production each have distinct signals upon which they activate^31^. Substantial work has shown that antibiotics contribute to CDAD by changing commensal structure and removing inhibition on both germination and outgrowth^32^. Yet, the specific structural definition of a CDAD-inhibitory gut microbiome remains elusive.

Our data suggest a potential mechanism by which an already outgrown but microbiome-controlled population of *C. difficile* might be able to take hold and produce toxin following the host changes of CDAD-associated antibiotics: loss of mucin barrier, increased sensitivity to toxin, and reduced innate immune response. We found that both CDAD-associated antibiotics lead to increased toxin transport (Fig. 2D) and concomitant sensitivity to toxin B (Fig. 4C, D).

Increased relative abundance of proteobacteria is associated with CDAD and has been proposed to be a risk factor^33^. Proteobacterial bloom following antibiotic treatment might be accounted for by the loss of *E. coli* and LPS responsiveness we uncovered.

Host-directed therapies might circumvent recurrent or drug resistant infection or prevent CDAD completely. Host response is a better predictor of patient outcome than specific changes to the microbiome or even than *C. difficile* bacterial load^34^, suggesting host stratification might be effective in preventing CDAD. Future work is required to identify potential host-directed therapies that might increase mucin production or innate immunity in the colon of patients taking high risk antibiotics. This work suggests using caution when prescribing CDAD-associated antibiotics, particularly to those at higher risk for CDAD.

Our work defines a new role for effects of CDAD-associated antibiotics on CDAD pathology, namely, the commensal-independent effects on the host. By reducing barrier function and immune cell capability and increasing toxin sensitivity, antibiotics with high risk for CDAD may prime the host to be less prepared for combating *C. difficile* infection and pathogenesis. This has important implications for potential host-directed prophylactic or CDAD-treatment therapies. Further work is needed to understand the commensal-independent effects of other antibiotics that might similarly prime the gut for enteric infection.

## Methods

### Tissue culture: cell lines

Caco2 (clone: C2BBe1, passage 48–58, ATCC, Manassas, VA) and HT29-MTX (passage 20–30, Sigma–Aldrich, St. Louis, MO) were maintained in DMEM (Gibco, Gaithersburg, MD) supplemented with 10% heat-inactivated FBS (Atlanta Biologicals, Flowery Branch, GA), 1% GlutaMax (Gibco), 1% Non-Essential Amino Acids (NEAA, Gibco), and 1% Penicillin/Streptomycin (P/S). Both cell lines were passaged twice post thawing before their use for transwell seeding. Briefly, the apical side of transwell membrane were coated with 50mg/mL Collagen Type I (Corning Inc., Corning, NY) overnight at 4°C. Caco2 at 80–90% confluence and HT29-MTX at 90-95% confluence were harvested using 0.25% Trypsin-EDTA and mechanically broken up into single cells. 9:1 ratio of C2BBe1 to HT29-MTX was seeded onto 12-well 0.4mm pore polyester transwell inserts (Corning, Tewksbury, MA) at a density of 10^5^ cells/cm^2^. Seeding media contained 10% heat inactivated FBS, 1x GlutaMax and 1% P/S in Advanced DMEM (Gibco). Seven days post seeding, the media was switched to a serum-free gut medium by replacing FBS with Insulin-Transferrin-Sodium Selenite (ITS, Roche, Indianapolis, IN) and the epithelial cultures were matured for another 2 weeks. P/S was left out of media during experimental procedures.

### Tissue culture: primary cells

Colon organoids (enteroids) used in this study were established and maintained as previously described^35,36^. Endoscopic tissue biopsies were collected from the ascending colon of de-identified individuals at either Boston Children’s Hospital or Massachusetts General Hospital upon the donors informed consent. Methods were carried out in accordance to the Institutional Review Board of Boston Children’s Hospital (protocol number IRB-P00000529) and the Koch Institute Institutional Review Board Committee as well as the Massachusetts Institute of Technology Committee on the Use of Humans as Experimental Subjects. Tissue was digested in 2 mg ml^−1^ collagenase I (StemCell, cat. no. 07416) for 40 min at 37 °C followed by mechanical dissociation, and isolated crypts were resuspended in growth factor-reduced Matrigel (Becton Dickinson, cat. no. 356237) and polymerized at 37 °C. Organoids were grown in expansion medium (EM) consisting of Advanced DMEM/F12 supplemented with L-WRN conditioned medium (65% vol/vol, ATCC, cat. no. CRL-3276), 2 mM Glutamax (Thermo Fisher, cat. no. 35050-061), 10 mM HEPES (Thermo Fisher, cat. no. 15630-080), Penicillin/Streptomycin (Pen/Strep) (Thermo Fisher, cat. no. 15070063), 50 ng ml^−1^ murine EGF (Thermo Fisher, cat. no. PMG8041), N2 supplement (Thermo Fisher, cat. no. 17502-048), B-27 Supplement (Thermo Fisher, cat. no. 17502-044), 10 nM human [Leu15]-gastrin I (Sigma, cat. no. G9145), 500 μM N-acetyl cysteine (Sigma, cat. no. A9165-5G), 10 mM nicotinamide (Sigma, cat. no. N0636), 10 |iM Y27632 (Peprotech, cat. no. 1293823), 500 nM A83-01 (Tocris, cat. no. 2939), 10 μM SB202190 (Peprotech, cat. no. 1523072) 5 nM prostaglandin E2 (StemCell cat. no. 72192) at 37°C and 5% CO_2_. Organoids were passaged every 7 days by incubating in Cell Recovery Solution (Corning, cat. no. 354253) for 40 min at 4 °C, followed by mechanical dissociation and reconstitution in fresh Matrigel at a 1:4 ratio.

For 2D enteroid studies, at day 7 post passaging, colon organoids were collected, Matrigel was dissolved with Cell Recovery Solution for 40 min at 4 °C followed by incubation of Matrigel-free organoids in Trypsin (Sigma, cat. no. T4549) at 37 °C for 5 minutes. Organoids were mechanically dissociated into single cells, resuspended in EM without nicotinamide and 2.5 uM thiazovivin (Tocris, cat. no. 3845) in the place of Y27632, and seeded onto 24-well 0.4 μm pore polyester transwell inserts (Corning, 3493) coated with a 200 μg/mL type 1 collagen and 1% Matrigel mixture at a density of 1 × 10^5^ cells/transwell. After 3-4 days of incubation, monolayers were confluent and differentiation was initiated. For differentiation apical media was replaced with Advanced DMEM/F12 plus HEPES, glutamax, and Pen/Strep and basal media with differentiation medium (DM), which is EM without L-WRN conditioned medium, nicotinamide, prostaglandin E2 and Y27632, but supplemented with 100 ng ml^−1^ human recombinant noggin (Peprotech, cat. no. 120-10C) and 20% R-spondin conditioned medium (Sigma, cat. no. SCC111). Transepithelial electrical resistance (TEER) measurements were performed using the EndOhm-12 chamber with an EVOM2 meter (World Precision Instruments). At day 8 post seeding, the 2D enteroids were washed to remove P/S and used for further experimentation.

Monocyte-derived dendritic cells were used as the immune component of the gut when indicated. Briefly, peripheral blood mononuclear cells (PBMCs) were processed from Leukopak (STEMCELL Technologies, Vancouver, BC, Canada). Monocytes were isolated from PBMCs using the EasySep Human Monocyte Enrichment Kit (STEMCELL Technologies, 19058) and were differentiated in RPMI medium (Gibco) supplemented with 10% heat-inactivated FBS (Gibco), 50 ng/mL GM-CSF (Biolegend, San Diego, CA), 35 ng/mL IL4 (Biolegend), and 10 nM Retinoic acid (Sigma). After 7 days of differentiation (at day 19–20 post epithelial cell seeding), immature dendritic cells were harvested using PBS (Gibco) and seeded onto the basal side of the gut transwells in the absence of P/S 1 day prior to start of experiment. Macrophages were derived similarly, but with M-CSF (Biolegend) at 500 ng/mL.

### RNA preparation, qPCR

RNA was prepared using PureLink RNA Mini Kit (Invitrogen, Carlsbad, CA) according to manufacturer’s instructions. DNA removal was done on column with PureLink DNase (Invitrogen). cDNA synthesis using High-Capacity RNA-to-cDNA Kit (Applied Biosystems, Foster City, CA) according to product insert. qPCR was completed using TaqMan^®^ assays with Fast Advanced Master Mix (Applied Biosystems) as per manufacturer’s guidelines.

### 3’ DGE library preparation

RNA samples were quantified and quality assessed using an Advanced Analytical Fragment Analyzer. 20ng of totalRNA was used for library preparation with ERCC Spike-in control Mix A (Ambion 10-6 final dilution). All steps were performed on a Tecan EVO150. 3’DGE-custom primers 3V6NEXT-bmc#1-24 are added to a final concentration of 1.2uM. (5’-/5Biosg/ACACTCTTTCCCTACACGACGCTCTTCCGATCT[BC_6_]N_10_T_30_VN-3’ where 5Biosg = 5’ biotin, [BC6] = 6bp barcode specific to each sample/well, N10 = Unique Molecular Identifiers, Integrated DNA technologies). After addition of the oligonucleotides, samples were denatured at 72C for 2 minutes followed by addition of SMARTScribe RT per manufacturer’s recommendations with Template-Switching oligo5V6NEXT (12uM, [5V6NEXT: 5’-iCiGiCACACTCTTTCCCTACACGACGCrGrGrG-3’ where iC: iso-dC, iG: iso-dG, rG: RNA G]) and incubation at 42C for 90’ followed by inactivation at 72C for 10’. Following the template switching reaction, cDNA from 24 wells containing unique well identifiers were pooled together and cleaned using RNA Ampure beads at 1.0X. cDNA was eluted with 90 ul of water followed by digestion with Exonuclease I at 37C for 45 minutes, inactivation at 80C for 20 minutes. Single stranded cDNA was then cleaned using RNA Ampure beads at 1.0X and eluted in 50ul of water. Second strand synthesis and PCR amplification was done using the Advantage 2 Polymerase Mix (Clontech) and the SINGV6 primer (10 pmol, Integrated DNA Technologies 5’-/5Biosg/ACACTCTTTCCCTACACGACGC-3’). 12 cycles of PCR was performed followed by clean up using regular SPRI beads at 1.0X, and was eluted with 20ul of EB. Successful amplification of cDNA was confirmed using the Fragment Analyzer. Illumina libraries are then produced using standard Nextera tagmentation substituting P5NEXTPT5-bmc primer (25μM, Integrated DNA Technologies, (5’-AATGATACGGCGACCACCGAGATCTACACTCTTTCCCTACACGACGCTCTTCCG*A*T *C*T*-3’ where * = phosphorothioate bonds.) in place of the normal N500 primer.

Final libraries were cleaned using SPRI beads at 1X and quantified using the Fragment Analyzer and qPCR before being loaded for paired-end sequencing using the Illumina NextSeq500.

### Sequencing data analysis

Post-sequencing, quality-control on each of the libraries was performed to assess coverage depth, enrichment for messenger RNA (exon/intron and exon/intergenic density ratios), fraction of rRNA reads and number of detected genes using bespoke scripts. The sequencing reads were mapped to hg38 reference using star/2.5.3a. Gene expression counts were further estimated using ESAT v1^37^.

### Data availability statement

The data discussed in this publication have been deposited in NCBI’s Gene Expression Omnibus (Edgar et al., 2002) and are accessible through GEO Series accession number GSE135383 (https://www.ncbi.nlm.nih.gov/geo/query/acc.cgi?acc=GSE135383).

### Phagocytosis and Intracellular Killing assays

*Escherichia coli* GFP (ATCC^®^ 25922GFP™) (*E. coli*) was grown to early log in LB media plus ampicillin, then washed and resuspended in RPMI without antibiotic at the appropriate density. PBMC-derived macrophages were treated with low dose (Supplemental Table 1) of indicated antibiotic for 3 days in RPMI with heat-inactivated FBS (Atlanta Biologicals). Antibiotic was removed and antibiotic-free RPMI with GFP+ *E. coli* was added at an MOI of 10:1. Extracellular *E. coli* were washed off at indicated time. For phagocytosis, at 30 minutes post infection, cells were fixed and permeabilized, DAPI (4’,6-diamidino-2-phenylindole, Thermo Fisher) and ActinRed™ 555 ReadyProbes™ (Molecular Probes, Life Technologies, Carlsbad, CA) stained, and imaged. Percent macrophages with at least one GFP+ bacterium was calculated from fluorescent microscopy images. For intracellular survival assay, macrophages were lysed in water, and supernatants were plated for CFU.

### Self-organizing map

Gene expression fold changes from controls across antibiotics were considered as a function of CDAD risk and were normalized to be between 0 and 1 across the 3 antibiotics used. Genes without expression fold changes across all 3 conditions were omitted. The map was initialized with a 2-dimensional 3×3 square grid and implemented using the MATLAB (MathWorks, Natick, MA) R2017b Neural Network Toolbox.

### Gene Ontology Enrichment Analysis

Gene ontology enrichment on all GO terms was performed using the free online PANTHER overrepresentation test^38–40^. FDR was set to <0.05.

### Data Representation and Statistical Analysis

Prism 8 software (GraphPad Software, La Jolla, CA) was used to graph all data except SOMs. Statistical tests of measurements were used from the Prism suite as noted in figure legends. Statistical significance is indicated by as follows: *<0.05, **<0.01, ***<0.001, ****<0.0001.

### Viability/Cytotoxicity analysis

Viability of monolayers post-antibiotic treatment was assessed using the Viability/Cytotoxicity Assay for Animal Live & Dead Cells kit (Biotium, Fremont, CA) according to package insert. Ratio of red to green cells was measured using ImageJ.

### Soluble mucin quantification by Alcian blue colorimetric assay

Apical medium was stained for 2 h at room temperature with 1% Alcian blue (Electron Microscopy Sciences) at a ratio of 1:4 with the sample. Dye-treated mucin was sedimented by a 30-min centrifugation (), followed by two wash steps in wash buffer (290 mL 70% ethanol, 210 mL 0.1 M acetic acid, and 1.2 g MgCl_2_). Dye-treated mucin was resuspended in 10% SDS and absorbance was read on a plate reader at 620 nm. Calculations were made based on a known standard prepared in parallel.

### Western blotting

Western blotting was performed under reducing conditions using iBlot 2 dry blotting system (Invitrogen) standard procedures. Primary antibodies were incubated at 4°C overnight diluted as noted in Odyssey blocking buffer: rabbit monoclonal anti-Vinculin antibody (abcam, Cambridge, MA [EPR8185]) at 1:3000, mouse monoclonal anti-Rac1 antibody (abcam, [23A8]) at 1:750, and purified mouse Anti-Rac1 antibody (BD Transduction Laboratories, San Jose, CA [clone 102/Rac1]) at 1:750. For detection, LI-COR (Lincoln, NE) goat anti-rabbit or anti-mouse IR800-conjugated secondaries at 1:8,000 were incubated for 30 minutes at room temperature in Odyssey blocking buffer (TBS) with 0.1% Tween 20. Imaging of membrane using LI-COR Odyssey imager with settings as follows: 24 μm resolution and high quality, laser intensity of 2.0 on the 800 channel.

### Chemokine quantification

Secreted IL-8 was measured by Quantikine^®^ ELISA Human IL-8/CXCL8Immunoassay (R&D Systems, Minneapolis, MN) per manufacturer’s guidelines.

## Acknowledgements

JCK supported by an administrative supplement to NIH R01-EB021908. DKB supported by Research Beyond Borders Program of Boehringer Ingelheim Pharmaceuticals. This work was supported in part by the Koch Institute Support (core) Grant P30-CA14501 from the National Cancer Institute and NIEHS Grant P30-ES002109. We thank the members of the LGG and DAL labs for their scientific input.

## Author Contributions Statement

JCK and LGG conceived of the project. JCK designed the experiments. JCK acquired and analyzed the data, with technical support from JV and CW. JCK and DKB performed self-organizing maps analysis, with critical validation from DAL. JCK drafted the original manuscript. All authors contributed to manuscript revisions. LGG and DAL provided oversight and leadership for the project. LGG provided grant support for all study materials and reagents.

